# Stationary-Phase *Pseudomonas aeruginosa* Fluoroquinolone Persisters Mostly Avoid DNA Double-Stranded Breaks

**DOI:** 10.1101/2025.08.29.673110

**Authors:** Patricia J. Hare, Juliet R. González, Wendy W.K. Mok

## Abstract

When susceptible bacterial cultures are treated with antibiotics, some cells can survive treatment without heritable resistance, giving rise to susceptible daughter cells in a phenomenon termed antibiotic persistence. Current models of fluoroquinolone (FQ) persistence in stationary-phase cultures posit that post-treatment resuscitation is dependent on double-stranded break (DSB) repair through RecA-mediated homology-directed repair. Previously, we found that stationary-phase *P. aeruginosa* does not depend on RecA to persist. In this work, we ask whether *P. aeruginosa* FQ persisters from stationary-phase cultures require DSB repair at all. We measured DSB formation in Levofloxacin (LVX)-treated cells recovering from treatment using strains expressing fluorescently labeled DSB-binding protein, Gam. We find that, surprisingly, the majority of *P. aeruginosa* LVX persisters survive treatment without apparent DSBs. Persisters that have evidence of DSBs take longer until their first division compared to persisters without DSBs, and the phenotypes of their progeny suggest how persisters cope with DSBs—via repair or by damage sequestration—in order to successfully propagate. These observations pave the way for mechanistic studies into *P. aeruginosa* FQ persistence and highlight the need for single-cell tools to track FQ-induced damage.

## IMPORTANCE

*Pseudomonas aeruginosa* is an opportunistic pathogen of significant clinical interest. When susceptible cultures of *P. aeruginosa* are treated with fluoroquinolone (FQ) antibiotics, some cells survive treatment and regrow in a phenomenon termed antibiotic persistence. Studies in *Escherichia coli* and other bacterial species suggest that FQ persisters survive by repairing DNA double-stranded breaks (DSBs) after antibiotic removal. In this study, we show that most stationary-phase *P. aeruginosa* survive by avoiding DSBs rather than repairing them.

## OBSERVATION

The fluoroquinolones (FQ) are a synthetic class of antibiotics that trap bacterial topoisomerases on DNA, leading to mounting topological stress and DNA double-stranded breaks (DSBs) (1, 2). Even in clonal laboratory cultures of FQ-susceptible bacterial populations, some cells can survive treatment in the phenomenon known as persistence. After treatment, persisters can lead to population resurgence and treatment failure (3, 4). Studies on FQ antibiotic persistence in stationary-phase *Escherichia coli, Staphylococcus aureus*, and *Salmonella spp*. have elucidated some commonalities across all species: i) the majority of cells that die do so in the post-antibiotic period, not during treatment itself, and ii) persisters engage in RecA-mediated DSB repair after FQ treatment to survive (5-8).

We previously found that FQ-treated stationary-phase *Pseudomonas aeruginosa* does not adhere to the first paradigm and dies readily during exposure to the FQ, Levofloxacin (LVX) (9). We also found that *P. aeruginosa* does not require RecA to persist after LVX treatment (9). Therefore, we hypothesized that *P. aeruginosa* also subverts the second tenet of FQ persistence and that this gram-negative species does not engage in DNA repair in order to persist.

To test this hypothesis, we modified a Gam reporter construct that has previously been utilized for labeling DSBs in *E. coli* and eukaryotic cell cultures (10, 11). We expressed IPTG-inducible Gam-mScarlet from the *P. aeruginosa* chromosome. Cells were induced to express Gam-mScarlet throughout growth to stationary phase then treated with LVX; after treatment, cells were seeded onto nutritive agarose pads for imaging during recovery. The appearance of fluorescent foci within cells indicates Gam aggregation onto dsDNA ends (**Fig. 1a**). Although Gam has been reported to obstruct the RecBCD nuclease and inhibit homologous recombination (HR) in *E. coli*, we did not find any differences in FQ persistence for *P. aeruginosa* strains with or without the fluorescent Gam construct (**Fig. 1b**) (12).

**Figure 1.**
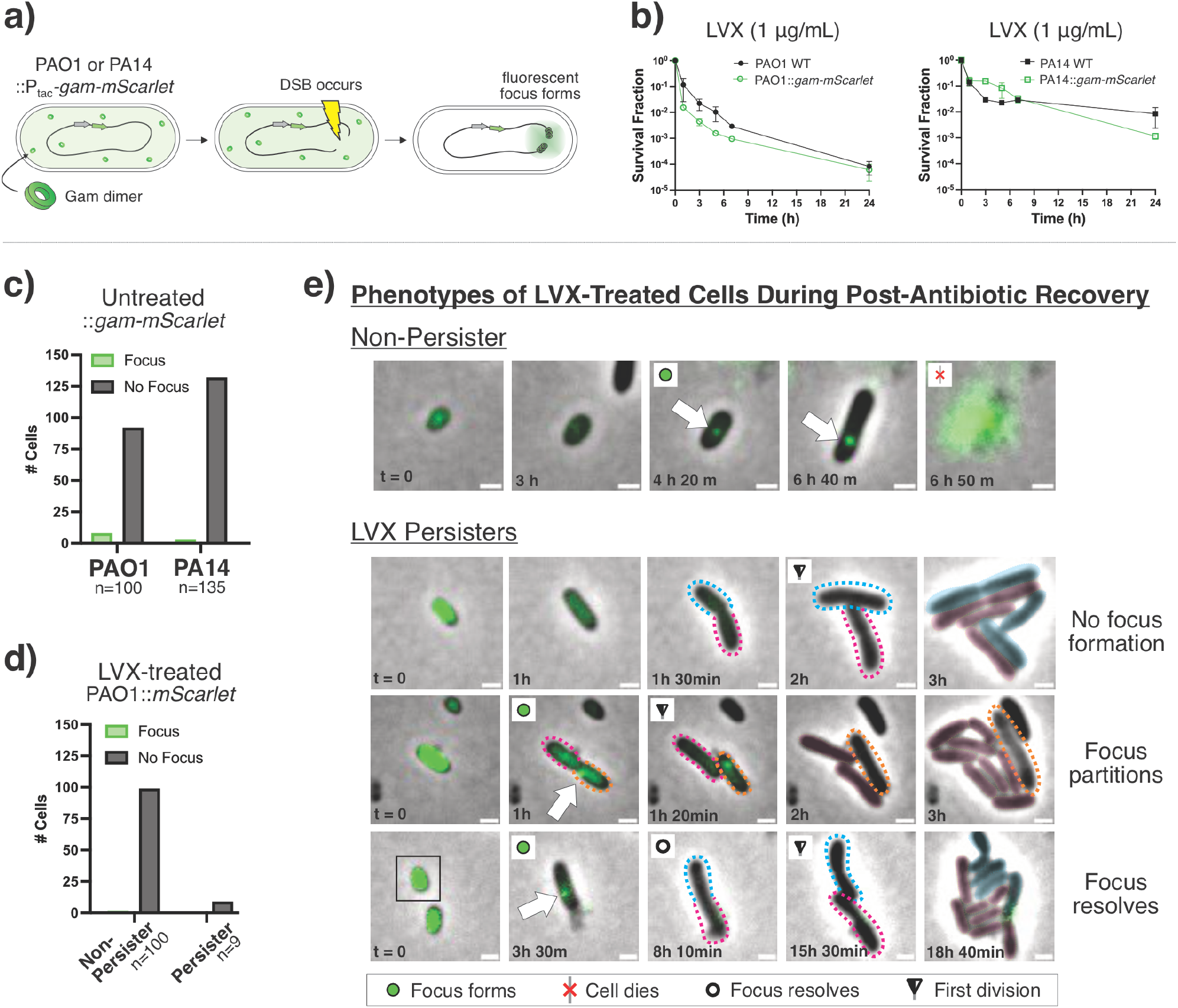
Fluorescently labeled Gam highlights DSBs in individual LVX-treated *P. aeruginosa* cells. **a)** *P. aeruginosa* strains bearing IPTG-inducible Gam-mScarlet were grown to stationary phase and treated with LVX (1 μg/mL). Fluorescent focus formation was tracked throughout post-antibiotic recovery. mScarlet fluorescence is false-colored as green for clarity. **b)** The survival of *P. aeruginosa gam-mScarlet* strains treated with 1 μg/mL LVX is not significantly different from wild-type strains. Fluorescent foci do not form in **c)** untreated *gam-mScarlet* strains or in **d)** PAO1::P_tac_-*mScarlet*—that lacks the Gam protein—during recovery from LVX treatment. **e)** Representative images of *P. aeruginosa gam-mScarlet* cell fates. The colored, dotted outlines demarcate daughter cells from the same persister; the corresponding color masks in later frames indicate the lineage from which progenies were derived. The orange outline indicates a non-dividing daughter cell. Scale bars represent 1 μm. Representative videos of PAO1 and PA14 with each phenotype can be found in the **Supplemental Materials**.

We verified the specificity of the fluorescent Gam assay by analyzing untreated Gam-mScarlet cells for both PAO1 and PA14 and saw that they rarely form fluorescent foci (**Fig. 1c**). This indicates that focus formation is not likely due to non-specific aggregation of Gam-mScarlet. We also treated cells bearing IPTG-inducible mScarlet with LVX and did not observe focus formation during recovery (**Fig. 1d**). From these controls, we conclude that foci formation is not attributable to non-specific mScarlet aggregation upon FQ treatment.

### FQ-treated cell phenotypes are heterogeneous during recovery

As Gam-mScarlet strains recovered from LVX treatment, we observed focus formation in dead or non-dividing cells (“non-persisters”) and persisters alike (**Fig. 1e**). For persisters that formed foci, we observed that foci either partitioned into one of the daughter cells or resolved before division (**Fig. 1e**). We analyzed imaging data for individual cells over 24 h recovery and plotted the times of focus formation (if any), focus resolution, cell death, and first cell division (persisters only) (**Fig. 2a, c**). Analyses were carried out for a random sample of at least 100 non-persister cells for each experimental replicate and all the persister cells in the given field(s) of view.

**Figure 2.**
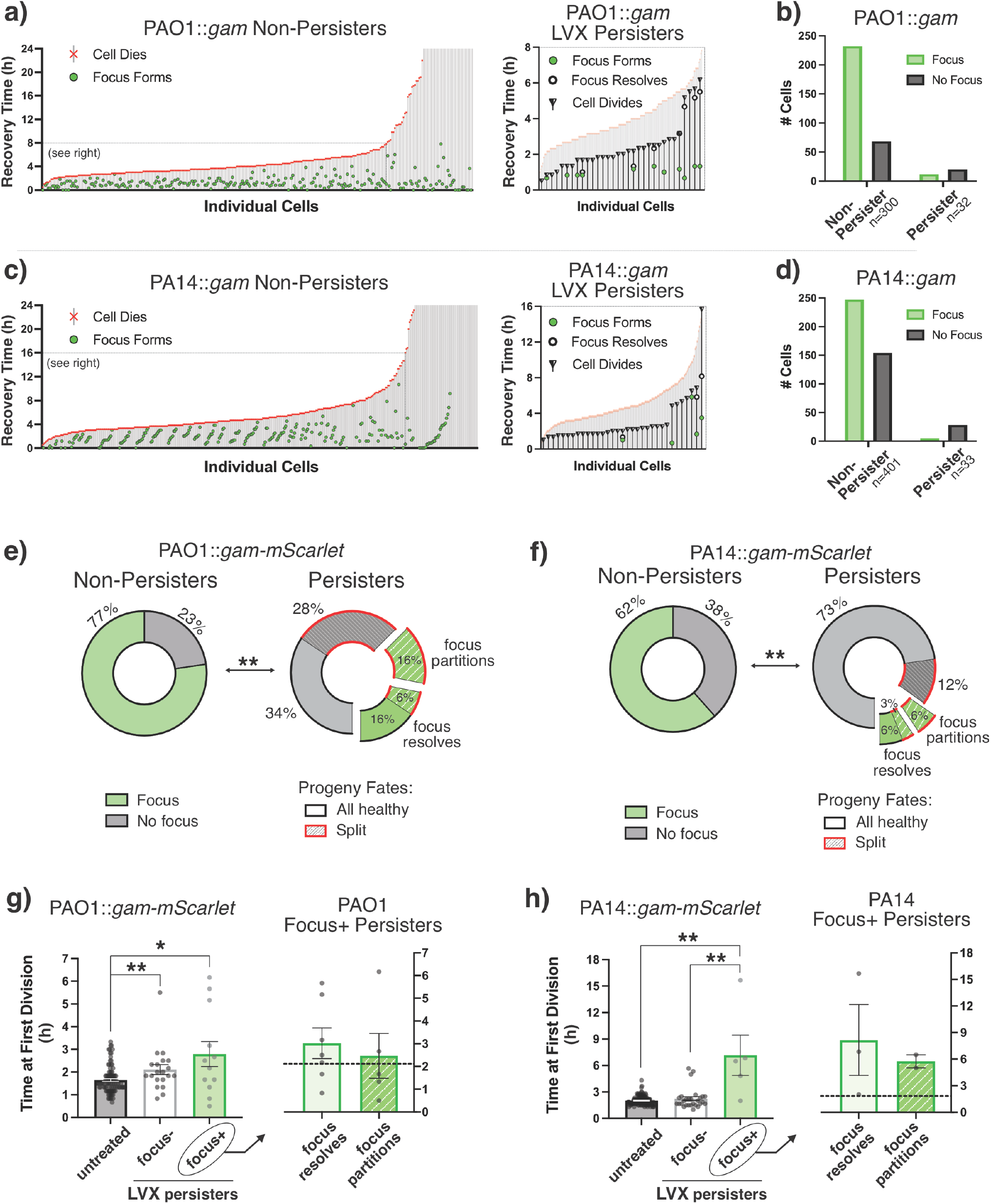
Imaging of *P. aeruginosa* Gam Strains Illustrates that Most LVX Persisters Avoid DSBs. The fates of individual *P. aeruginosa*::*gam-mScarlet* cells were tracked during recovery from LVX treatment (1 μg/mL) in **a)** PAO1 and **c)** PA14 (n > 300 non-persisters over > 2 biological replicates per strain). Cells that never divided or died after 24 h recovery were categorized as non-persisters. To the right, fates for persisters are shown overlaid onto the non-persisters. **b, d)** Aggregate data show that untreated cells and persisters rarely form fluorescent foci, but non-persisters do. **e, f)** The ratios of fluorescent: non-fluorescent cells are significantly different between non-persisters and persisters for both strains (** denotes p < 0.01; two-sided Fisher’s exact test). The pie charts show the fates of persisters that did or did not form foci and whether their progeny were healthy (daughter cells successfully divide) or split (one daughter cell fails to divide/lyses). **g, h)** The times of initial cell division are shown for each persister focus phenotype compared to untreated Gam-mScarlet strains. Statistical significance was assessed using unpaired, two-tailed Mann-Whitney tests with Welch’s correction for unequal variance (* indicates p < 0.05; ** p < 0.01). The graph to the right shows time of division for persisters that formed foci; the dashed line represents the average time to first division for non-focus forming persisters.

The visual summaries of individual cell fates show that there is widespread post-antibiotic killing of *P. aeruginosa* treated with LVX (**Fig. 2a, c**). Most non-persisters died within the first 4.5 hours (median times of death: 4.17 h for PAO1, 4.5 h for PA14). Of note, a significantly greater proportion of non-persister cells formed fluorescent foci during recovery than persister cells (**Fig. 2b, d-f**). This suggests that *P. aeruginosa* persisters mostly do not form DSBs when recovering from LVX treatment.

### FQ persister progenies are a mix of viable and non-viable cells

We noticed that persisters either divided into healthy, proliferative daughter cells or the progeny were split between viable and non-dividing cells (13). Regardless of whether a persister had Gam focus formation, cells were seen to give rise to both healthy and split progeny (**Fig. 2e, f**). For the persisters that formed Gam foci, the likelihood that foci dissipated (“resolved”) or were partitioned into one of the daughter cells at the time of division was comparable. Of the PAO1 and PA14 persisters that partitioned foci, all seven of them gave rise to split progeny (**Fig. 2e, f**). Note that the daughter cells that retained foci failed to proliferate, suggesting that segregation of the Gam-labeled DSB into one daughter cell allowed the other to persist.

### FQ persisters with DSBs have longer exit from lag

We observed that LVX persisters with foci formation took significantly longer to exit lag and divide than cells that were never treated with LVX (**Fig. 2g, h**). By comparison, the time of first division for foci-less persisters was similar to that of untreated cells. Persisters without foci divided approximately 40 min and 2 h earlier than the persisters with foci, respectively, for PAO1 and PA14. These data support a model in which persisters that avoid DSBs are able to resume growth quickly like untreated stationary-phase cells, outpacing persisters that must cope with DSBs before dividing.

## Discussion

Consistent with our previous data showing that RecA-mediated DSB repair is not necessary for stationary-phase *P. aeruginosa* FQ persisters, our microscopy data suggest that persisters are cells that avoid, rather than repair, DNA damage (9). We observed that LVX persisters infrequently form fluorescent Gam foci (indicative of DSBs) en route to propagating after FQ treatment (**Fig. 2a, c**). It is possible that FQ treatment might generate DSBs with single-stranded overhangs that cannot be bound by Gam (11). However, the frequency of fluorescent foci in non-persister cells suggests that Gam-detectable breaks *do* occur in the majority of recovering FQ-treated cells: 77% and 62% for PAO1 and PA14 non-persisters, respectively (**Fig. 2e, f**). The infrequency of Gam foci in persister cells seems to support the traditional perspective that *P. aeruginosa* FQ persisters are cells with low metabolic activity that are less susceptible to injury by antibiotics that target active cell processes (14, 15).

In keeping with this model, we found that persisters took more time to initially divide after treatment than untreated cells (**Fig. 2g, h**). Longer exit from lag is an established trait of metabolically quiescent cells that have increased persistence against antibiotics (16). Conversely, the delay may actually be a symptom, reflecting the time it takes for a persister to repair damage, liberate FQ-topoisomerase-DNA ternary complexes, or expel residual antibiotic before propagating (17, 18). We hypothesize that the mode of Gam focus resolution and fates of progeny indicate specific persistence mechanisms. For persisters whose fluorescent foci dissipated, we hypothesize that the longer delay until first division—compared to persisters without fluorescent foci—indicates the time it takes for break repair enzymes to displace Gam and complete repair. For persisters that partitioned foci into daughter cells, we hypothesize that those cells were effectively sorting FQ-trapped ternary complexes into their daughter cells, condemning some to die so that the others might propagate (17, 18). We expect that repair takes longer than ternary complex segregation and might explain those persisters’ slightly longer times until first division (**Fig. 2g, h**).

Collectively, our data suggest that *P. aeruginosa* FQ persisters do not fit the paradigms set by other pathogens. Furthermore, these data provide the impetus for further mechanistic studies of ternary complex segregation in FQ persister progeny. Understanding how individual cells overcome FQ treatment will inform strategies for fully eradicating susceptible populations, thereby limiting an infection’s ability to develop antibiotic resistance (4, 19, 20).

## Acknowledgements

We thank Dr. Keith Poole for distributing *P. aeruginosa* PAO1 (K767); Dr. Mona Wu Orr for distributing *E. coli* HB101 pRK2013, *E. coli* DH5α pJM252, and *E. coli* SM10 pFLP2; and Dr. Susan Rosenberg for sharing pRF3-Gam with us. We thank Dr. Yi Wu and Ms. Susan Staurovsky (UConn Center for Cell Analysis and Modeling Microscopy Facility) for their assistance with experiments. We are grateful to Dr. Barbara Kazmierczak and Dr. Rebecca Page for their insights and discussions.

This work was supported by funding awarded to W.K.M. from the University of Connecticut start-up fund and the National Institutes of Health (NIGMS; R01GM147257 and DP2GM146456). P.J.H. was additionally supported by the National Institutes of Health (NIDCR; F30DE032598) and previously by the NIH Skeletal, Craniofacial, and Oral Biology Training Grant (T90DE021989). The funders had no role in the design of our experiments or preparation of this manuscript.

