## Supplemental Materials for "Stationary-Phase *Pseudomonas aeruginosa* Fluoroquinolone Persisters Mostly Avoid DNA Double-Stranded Breaks"

Materials and Methods

Tables
- Table S1
- Table S2
- Table S3

Supplemental Video Captions
- Video S1
- Video S2
- Video S3
- Video S4
- Video S5
- Video S6
- Video S7
- Video S8
- Video S9
- Video S10

References

### Materials and Methods

#### Culture Media and Antibiotics

Cells were inoculated from frozen stocks into cation-adjusted Mueller-Hinton Broth (CA-MHB) and cultured for 4 h. CA-MHB was prepared from BD Difco Mueller Hinton Broth powder and cation-adjusted to final concentrations of 10 mg/L $Mg^{2+}$ and 20 mg/L $Ca^{2+}$; cations were prepared as 85 g/L $MgCl_2 \cdot 6H_2O$ and 28 g/L $CaCl_2$ stock solutions in water and filter-sterilized before adding to autoclaved MHB media. In order to decrease possible confounding factors due to batch variation of rich media, cultures for phenotypic assays were subcultured after the 4 h pre-growth into Basal Salt Media (BSM), a chemically defined minimal media with succinate as the sole carbon source (1, 2). BSM was prepared in water with 30.8 mM $K_2HPO_4$, 19.3 mM $KH_2PO_4$, 15 mM $(NH_4)_2SO_4$, 1 mM $MgCl_2$, 2 μM $FeSO_4$, and 15 mM succinic acid, then filter-sterilized before use. All liquid cultures were incubated at 37 °C, shaking at 250 rpm.

Antibiotic stocks were prepared at the following concentrations: Levofloxacin (LVX) - 5 mg/mL in water, Tetracycline (Tet) – 10 mg/mL in water, Igrasan (Igr) - 25 mg/mL stock in ethanol, Carbenicillin (Carb) – 100 mg/mL in water, Ampicillin (Amp) – 100mg/mL in water. Agar plates were prepared as 1.5% agar. Antibiotics prepared in water were filter-sterilized using 0.22 μm polyethersulfone (PES) filters before use.

#### Preparation of the miniCTX-Gam-mScarlet Integrative Vector

A gene fragment containing $P_{lacIQ}$-*lacI*-$P_{tac}$-mScarlet-I was ordered (Integrated DNA Technologies) with restriction sites on either end and amplified using lacIQ_Ptac_FWD2 and mScarlet_REV primers (see **Table S3**). The amplified gene was ligated into pJM252 by restriction digest at the KpnI and SacI cut sites. The ligation products were transformed into chemically competent *E. coli* DH5α and transformants were selected on Tet10 agar plates. Plasmids were confirmed by whole plasmid sequencing. The resultant miniCTX-Ptac-mScarlet vector was used as the backbone for creating the Gam-mScarlet translational fusion vector.

Gam was amplified from pRF3-Gam using primers Gam_FWD and Gam_REV, which contained restriction sites for SacI and SpeI, respectively (3). The Gam_REV primer also included an extra four alanine residues to create a flexible linker between Gam and mScarlet. The amplified gene was ligated into miniCTX-$P_{tac}$-mScarlet by restriction digest at the SacI and SpeI cut sites. The ligation products were transformed into chemically competent *E. coli* DH5α and cells were selected on Tet10 agar plates. Plasmids were confirmed by whole plasmid sequencing.

#### Cloning of Insertion Strains Using miniCTX Vector Derivatives

*P. aeruginosa* Gam-mScarlet and mScarlet strains were generated using the miniCTX vector which integrates genes of interest into the neutral *attB* site of the *P. aeruginosa* chromosome (4). The *P. aeruginosa* parental strain ("recipient") was transformed via conjugation by triparental mating with *E. coli* containing the miniCTX derivative ("donor") and the helper strain *E. coli* HB101 pRK2013.

After triparental mating, transformed *P. aeruginosa* were selected for on LB agar containing Igr (25 μg/mL) and Tet (75 μg/mL). Three colonies on the LB-Igr25-Tet75 plate were grown in plain

LB for 2-3 h to cure miniCTX, then each clone was plated onto its own antibiotic-free LB agar plate to create a lawn. Simultaneously, lawns of *E. coli* SM10 + pFLP2 were plated onto LB + Carb (300 μg/mL) agar plates. The next day, pFLP2 was transformed into each *P. aeruginosa* mutant by biparental mating on plain LB agar. After 2 h mating, the mating spot was scraped, resuspended in LB, and dilutions were plated to LB agar and LB + 10% sucrose agar. The next day, sucrose-resistant clones were patched onto 1) plain LB agar, 2) LB + 10% sucrose agar, 3) LB + Carb300 agar, and 4) LB + Tet75 agar. Clones which were sucrose-resistant (indicates loss or inactivation of *sacB* gene on pFLP2), Tet-sensitive (indicates Tet resistance marker was cured from the miniCTX insert), and Carb-sensitive (indicates cured of pFLP2, a back-up to the 10% sucrose plate) were picked from the plain LB plate, grown in plain LB, and screened by cPCR for the insert of interest.

**Whole Genome Sequencing**

Genomic DNA was collected from individual clones grown in LB using the Qiagen Blood & Tissue Kit, according to the manufacturer's protocols. DNA was sent to SeqCenter (Pittsburgh, PA) for library preparation and sequencing. Libraries were prepared using the Illumina DNA Prep kit with custom 10 bp unique dual indices (UDI) with a target size of 280 bp. Libraries were sequenced on an Illumina NovaSeq X Plus (150 bp paired end reads, 2.67 million reads per sample). The resultant fastq read files were submitted to the online genome visualization tool, Proksee (5), for genome assembly (Proksee Assemble version 1.0.0a6) and feature annotation via Prokka (version 1.14.6) (6). To verify specific sequences, contigs with the regions of interest were downloaded from Proksee and compared to the desired sequences using SnapGene Viewer (version 7.0.1). Raw WGS reads can be found on the National Center for Biotechnology Information (NCBI) Sequence Read Archive under BioProject PRJNA1288492.

**LVX Persistence Assays**

*P. aeruginosa* strains were inoculated from -80 °C frozen stocks into 2 mL test tubes of CA-MHB and grown at 37 °C, shaking at 250 rpm. After 4-5 h of growth, inoculations were diluted 1:100 into 250-mL baffled flasks with 25 mL BSM. Following growth to stationary phase (16 h), $OD_{600}$ of each culture was measured and 10 μL cells were collected for serial dilution in PBS and plating onto CA-MHB agar plates for colony forming unit (CFU) enumeration. Cells were then treated with 1 μg/mL LVX. At designated timepoints, 500 μL culture was collected and pelleted by centrifugation at 21,000 x g for 3 min. After removing 450 μL of supernatant, the pellets were washed with 450 μL PBS. This step was repeated, effectively diluting the antibiotics to subinhibitory levels (100-fold dilution). The cells were then serially diluted in PBS and 10 μL spots of each dilution were plated onto CA-MHB agar for CFU enumeration.

**Gam-mScarlet Assays and Time-Lapse Image Analysis of Gam Foci Formation**

PAO1 and PA14 bearing the Gam-mScarlet translational fusion or the $P_{tac}$-*mScarlet* control were grown for 16 h to stationary phase in BSM. At t=0, the $OD_{600}$ of each culture was measured and 10 μL cells were collected for serial dilution and plating onto CA-MHB agar for CFU enumeration. Then, the cultures were treated for 5 h with LVX (1 μg/mL). After treatment, cells were washed twice in PBS, and serially diluted for plating to CA-MHB for CFU enumeration. The washed cells were also diluted 30-fold in PBS for imaging. The cell dilutions were seeded onto antibiotic-free agarose pads made with BSM in a Bioptechs interchangeable cover dish (7). Cells were imaged every 10 min for 24 h recovery in a PeCon live cell incubation chamber kept at 37 $^{o}$C.

Images were analyzed using Fiji (ImageJ2 version 2.9.0/1.53t) (8). Phase channel image stacks were merged with the OFP channel (for mScarlet-I fluorescence) then the merged stacks were drift-corrected using the Correct_3D_Drift.py script (9, 10).

Cell tracking and morphological/fluorescent signal classification were conducted with the MicrobeJ plugin (version 5.13l) (11). The scale was set to 0.1032 μm/pixel. In brief, *P. aeruginosa* bacterial cells were detected on the Phase channel with the following parameters: Area (0.4 -10 μm), Length (0.1 - 20 μm), Width (0-2.5 μm), Curvature (0-0.7), Angularity (0-0.2), Exclude on Edges, Shape Descriptors, and Segmentation. The Tracking option with Lineage analysis was selected.

From the Results table, 100 numbers were randomly selected from the total number of bacteria in the first frame (t=0) using a random number generator. Cells that were already dead at the first imaging timepoint were not included in the analysis, as their focal fluorescence status could not be determined. Cells that were dead at frame 0 were excluded from the list of 100 cells and replaced by the next eligible listed cell. Cells that were persisters were also replaced by the next eligible listed cell. The 100 non-persister cells were tracked and the frames in which they formed a fluorescent focus (fluorescent signal $\geq$ 2x fluorescence of the local area in the cell) and/or died (by explosive lysis or shrinking lysis and loss of phase contrast) were recorded. For persisters or cells that had not been LVX-treated, the frame in which they first divided was also recorded. Analyses were conducted on images from at least two independent experimental replicates.

**Statistical Analysis**

All experiments were conducted over at least two biological replicates. Unless otherwise noted, statistical comparisons between samples were done by unpaired t-tests with Welch's correction for unequal variance at a significance level of 0.05.

**Table S1.** Bacterial strains used in this study.

| Strain (Genotype) | Description | Source |
|---|---|---|
| ***P. aeruginosa*** **PAO1** | wild-type | Poole Lab,<br>Queen's University<br>(12) |
| **PAO1::$P_{tac}$-*gam-mScarlet*** | IPTG-inducible<br>Gam-mScarlet-I fusion | This work |
| **PAO1::$P_{tac}$-*mScarlet*** | IPTG-inducible mScarlet-I | This work |
| ***P. aeruginosa*** **PA14** | wild-type<br>NR-50573 | BEI Resources |
| **PA14::$P_{tac}$-*gam-mScarlet*** | IPTG-inducible<br>Gam-mScarlet-I fusion | This work |
| ***E. coli*** **HB101**<br>**pRK2013** | F- *mcrB mrr hsdS*20(rB- mB-) *recA*13 *leuB*6 *ara*-14 *proA*2 *lacY*1 *galK*2 *xyl*-5 *mtl*-1 *rpsL*20(SmR) *glnV*44 λ-<br>mobilization helper plasmid; Kan$^R$ | Wu Orr Lab,<br>Amherst College |
| ***E. coli*** **DH5α** | F- 80d*lacZ* M15 (*lacZYA-argF*) U169 *recA*1 *endA*1*hsdR*17(rk-, mk+) *phoA*supE44 -*thi*-1 *gyrA*96 *relA*1 | Wu Orr Lab,<br>Amherst College |
| ***E. coli*** **SM10 (λpir)**<br>**pFLP2** | *thi thr leu tonA lacY supE recA*::RP4-2-TcR::Mu *KmR* λpir<br>FLP recombinase plasmid; Carb$^R$ | Wu Orr Lab,<br>Amherst College |

**Table S2.** Plasmids used in this study.

| **Plasmid** (selective marker) | **Description** | **Source** |
|---|---|---|
| **pJM252** (Tet) | miniCTX-$P_{tac}$-$P_{lacIQ}$-*lacI* Integrative vector that localizes to the neutral *attB* site in the *P. aeruginosa* chromosome | Wu Orr Lab, Amherst College |
| **pFLP2** (Carb) | FLPase-containing plasmid for curing FRT-flanked antibiotic resistance markers | Wu Orr Lab, Amherst College |
| **pRF3-Gam** (Amp) | Vector with Gam-GFP | Rosenberg Lab, Baylor College of Medicine (3) |
| **miniCTX-mScarlet** (Tet) | Integrative vector with $P_{lacIQ}$-*lacI* and $P_{tac}$-*mScarlet-I* | This work |
| **miniCTX-gam-mScarlet** (Tet) | Integrative vector with $P_{lacIQ}$-*lacI* and $P_{tac}$-*gam-mScarlet-I* | This work |

**Table S3.** PCR Primers used for this study.

| Primer | Sequence (5' to 3') | Use |
|---|---|---|
| **lacIQ_Ptac_FWD2** | TCATCATGAGCTCGGATCCCGCTAACTTACATTAATTGCG | Amplify $P_{lacIQ}$-*lacI*-$P_{tac}$-mScarlet-I |
| **mScarlet_REV** | GTAGTAGGGTACCCTCGAGAAGCTTCTTGTACAGCTCGTC | Amplify $P_{lacIQ}$-*lacI*-$P_{tac}$-mScarlet-I |
| **Gam_FWD** | TCATCATGAGCTCCGTATCACGAGGCC | Amplify Gam from pRF3 |
| **Gam_REV** | GTAGTAGACTAGTGGCGGCGGCGGCAATACCGGCTTCCTGTTCAA | Amplify Gam from pRF3 |

### Supplemental Video Captions

**Videos S1-S5:**
Time-lapse fluorescence microscopy videos of *P. aeruginosa* PAO1::$P_{tac}$-*gam-mScarlet* during recovery from LVX treatment (1 μg/mL). These images are representative of three fields of view over two independent experiments. mScarlet fluorescence has been false-colored green for visual contrast. In each video, the cell of interest is centered and/or indicated by a box.

**Video S1:**
**PAO1::*gam* persister with no fluorescent focus formation and healthy progeny.**

**Video S2:**
**PAO1::*gam* persister that does not form fluorescent foci and gives rise to split progeny.**

**Video S3:**
**PAO1::*gam* persister with a partitioned fluorescent focus and split progeny.**

**Video S4:**
**PAO1::*gam* persister that gives rise to split progeny after resolving fluorescent foci.**

**Video S5:**
**PAO1::*gam* persister that gives rise to healthy progeny after resolving fluorescent foci.**

**Videos S6-S10:**
Time-lapse fluorescence microscopy videos of *P. aeruginosa* PA14::$P_{tac}$-*gam-mScarlet* during recovery from LVX treatment (1 μg/mL). These images are representative of four fields of view over two independent experiments. mScarlet fluorescence has been false-colored green for visual contrast. In each video, the cell of interest is centered and/or indicated by a box.

**Video S1:**
**PA14::*gam* persister with no fluorescent focus formation and healthy progeny.**

**Video S2:**
**PA14::*gam* persister that does not form fluorescent foci and gives rise to split progeny.**

**Video S3:**
**PA14::*gam* persister with a partitioned fluorescent focus and split progeny.**

**Video S4:**
**PA14::*gam* persister that gives rise to split progeny after resolving fluorescent foci.**

**Video S5:**
**PA14::*gam* persister that gives rise to healthy progeny after resolving fluorescent foci.**
